# Social foraging extends associative odor-food memory expression in an automated learning assay for *Drosophila*

**DOI:** 10.1101/636399

**Authors:** Aarti Sehdev, Yunusa G. Mohammed, Cansu Tafrali, Paul Szyszka

## Abstract

Animals socially interact during foraging and share information about the quality and location of food sources. The mechanisms of social information transfer during foraging have been mostly studied at the behavioral level, and its underlying neural mechanisms are largely unknown. The fruit fly *Drosophila melanogaster* has become a model for studying the neural bases of social information transfer, as fruit flies show a rich repertoire of social behaviors and provide a well-developed genetic toolbox to monitor and manipulate neuronal activity. Social information transfer has already been characterized for fruit flies’ egg laying, mate choice, foraging and aversive associative learning, however the role of social information transfer on associative odor-food learning during foraging are unknown. Here we present an automated learning and memory assay for walking flies that allows studying the effect of group size on social interactions and on the formation and expression of associative odor-food memories. We found that both inter-fly attraction and the duration of odor-food memory expression increase with group size. We discuss possible behavioral and neural mechanisms of this social effect on odor-food memory expression. This study opens up opportunities to investigate how social interactions are relayed in the neural circuitry of learning and memory expression.

## INTRODUCTION

Vertebrates often forage in groups to get a more accurate estimate of the location and quality of resources (Giraldeau and Caraco, 2000; Templeton and Giraldeau, 1996; Valone, 1989; Ward and Zahavi, 1973). Insects also convey information about the location and quality about a food source through social interactions. For example, honey bees signal the direction and distance of food locations to other bees (Frisch, 1965), ants complement their individual memory of a route to food using trail pheromones left by scouts (Czaczkes et al., 2011), and stimulus enhancement and local enhancement at the food source improves foraging efficiency in bumble bees (Alem et al., 2016; Avarguès-Weber and Chittka, 2014; Leadbeater and Dawson, 2017; Worden and Papaj, 2005). These social effects on foraging have been mostly studied at the level of behavioral outcome, and the neural mechanisms of how social information transfer improves foraging are still unknown.

The fruit fly, *Drosophila melanogaster*, is a suitable model organism for studying the effects of social interactions on foraging at both the behavioral and the neuronal level. Fruit flies are gregarious (Lefranc et al., 2001; Navarro and del Solar, 1975) and demonstrate a rich repertoire of social behaviors that encompass communication about internal states (Suh et al., 2004), social information spread during odor avoidance (Ramdya et al., 2014), foraging (Abu et al., 2018; Durisko and Dukas, 2013; Golden and Dukas, 2014; Lihoreau et al., 2016; Tinette et al., 2004) and predator-induced egg-retention (Kacsoh et al., 2015). Moreover, fruit flies socially learn, and naïve flies copy mate-choices (Danchin et al., 2018; Germain et al., 2016; Mery et al., 2009) and oviposition site-choices from experienced conspecifics (Battesti et al., 2012; Sarin and Dukas, 2009), and they show increased aversive odor memory retrieval when in groups (Chabaud et al., 2009). However, it is still unknown whether social information transfer affects flies’ associative odor-food learning during foraging.

The mechanistic understanding of foraging in fruit flies is unparalleled, both in regard to the neural mechanisms of odor-guided search (Galizia, 2014; Haverkamp et al., 2018; Wilson, 2013) and feeding (Itskov and Ribeiro, 2013), and of associative odor-food learning (Burke et al., 2012; Huetteroth et al., 2015; Liu et al., 2012; Owald and Waddell, 2015; Schwaerzel et al., 2003; Tempel et al., 1983; Thum et al., 2007), making the fruit fly a good model for studying the neural mechanisms of social interactions during foraging.

Here we investigated whether fruit flies socially interact during foraging and whether group size affects associative odor-food memory expression. We developed an automated assay to study associative odor-food reward learning and memory in single flies and in groups of flies. We found that odor-food memory expression increased in strength and duration with increasing group size, and flies in small or large groups, but not in pairs, were attracted to each other. These data confirm that flies socially interact during foraging (Abu et al., 2018; Durisko and Dukas, 2013; Golden and Dukas, 2014; Lihoreau et al., 2016; Tinette et al., 2004). In addition, these data suggest that social interactions increase the efficiency of odor memory-guided food search.

## MATERIALS AND METHODS

### Animals

*Drosophila melanogaster* wild type S were raised on a standard food medium (100 mL contain 6.7 g fructose, 2.4 g dry yeast, 0.7 g agar, 2.1 g sugar beet syrup, 0.282 g ethyl paraben and 0.61 ml propionic acid). Flies were raised in a room with natural day light cycle, with an average temperature of 23.5 °C and 32% relative humidity. One to four days old flies were anesthetized with CO_2_ and female flies were collected. Flies were starved for 2-3 days to motivate them to search for food. Flies were starved in a fly vial with filter paper soaked in water.

### Learning and memory assay

To condition groups of flies, we used an automated rotating platform with four circular arenas (Figure 1A, B). The arenas were covered with a watch glass (7 cm diameter, 8 mm height in the center) which was coated on the inner side with Sigmacote (Sigma-Aldrich) to prevent flies from walking on the inside of the glass. The floor was made of a Teflon-coated fiberglass fabric (441.33 P, FIBERFLON, Konstanz). Pure odorants (ethyl acetate and 2,3-butanedione, Sigma-Aldrich) were stored in 20 ml vials (Schmidlin Labor and Service). The vials were mounted under the platform and the lid was pierced with a needle (Hypodermic-needle; 0.45 x25 mm, Sterican), allowing the odorant to diffuse through a hole (diameter: 5 mm) in the platform through the Teflon fabric and into the arena. Each arena had two odorant sources. One odorant was used as sucrose-paired conditioned stimulus (CS+) and the other odorant was used as unpaired conditioned stimulus (CS-). Each odorant was used equally often as CS+ and CS-. At the location of the CS+, 20 μl of oversaturated sucrose-ethanol solution was pipetted onto the fiberglass fabric and blow-dried for 20 minutes, producing a thin layer of pure sucrose on a round patch with a diameter of 10 mm. The position of the CS+ and CS-were always switched between the conditioning and the test (e.g. if CS+ was at the inside position during conditioning, it was at the outside position during the test, and in half of the experimental runs the CS+ was at the inside position during conditioning, and in the other half at the outside position). To change the floor between experimental phases, the platform was rotated underneath the arenas; the arenas themselves did not move. The angular rotation speed of the platform was 360°/25 s which corresponded to a speed of 2.6 cm/s in the center of the arena (the distance between center of the platform and the center of the arena is 10.25 cm). The conditioning apparatus was placed in an air suction hood in order to remove odorants. All experiments were performed in the dark to eliminate visual stimuli. The arena was back-illuminated with infrared light (850 nm, SOLAROX LED Strip), which is not visible to flies, and experiments were video recorded with an infrared sensitive camera (infrared Camera Module v2, Pi NoiR, connected to a Raspberry Pi 3, Model B V1.2) at 15 frames/s. The rotating motor was controlled via TTL pulses through the Raspberry Pi. The rotation of the platform and the video recordings were controlled with custom-written software in Python (Stefanie Neupert).

**Figure 1:**
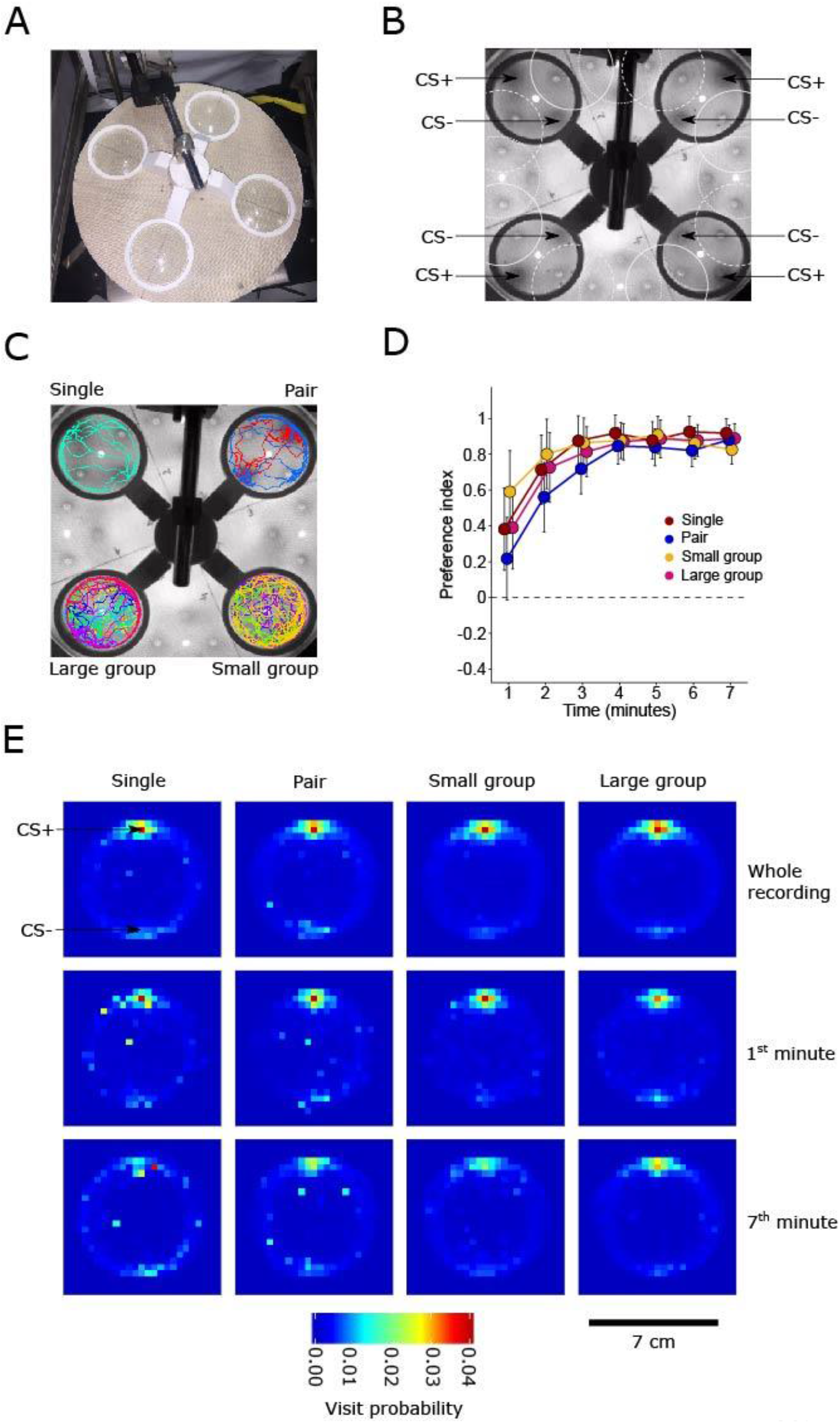
Automated odor-food learning and memory assay. (**A**) Odor-food learning and memory assay for four groups of flies. (**B**) Assay viewed from above through the tracking camera. The locations of the CS+ and CS− in each arena are indicated. The arcs indicate the position of the arena in different experimental phases. Full lines outline the acclimatization phase, dotted lines outline the conditioning phase, and dashed lines outline the pause phase before the test. (**C**) Same as (B) but with examples of individual fly trajectories during the test overlaid. Trajectories of a single fly (top left), a pair (top right), a group of 4 (small group, bottom right) and a group of 8 flies (large group, bottom left). (**D**) Preference for the odorant CS+-sucrose patch for the different groups during conditioning. Each minute bin shows the mean preference index across all experimental runs (N= 30 experimental runs). The dashed line represents the preference index of 0 (chance). In all time bins, all groups preferred the arena half containing the CS+-sucrose patch over the arena half containing the CS− (p(preference index > 0) ≥ 0.967). For statistical comparisons between groups see Table S1. (**E**) Visit probability maps for each group size (columns) during the memory test. Visit probabilities were calculated for the whole recording, the first minute of recording and the seventh minute of recording (rows). Each bin shows the mean binary value across all individuals of all 30 experimental runs (Single flies: N = 30 individuals; Pairs: N = 60 individuals; Small group: N = 118 individuals; Large group: N = 229 individuals).

### Experimental protocol

All experiments were done between 10:00 and 12:00 or after 15:00 during periods when flies show higher foraging activity (Breugel et al., 2017). Each experimental run contained 4 differently sized groups (“single”, “pair”, “small group”, “large group”), and the positions of the 4 arenas used for the 4 differently sized groups were balanced across experimental runs.

One experimental run consisted of four phases:

1. Acclimatization (Figure 1B, solid arcs): Flies were sucked out from the starvation vials using a tube aspirator and placed into an arena that had no odorant source and were allowed to acclimatize for 10 minutes.
2. Conditioning (Figure 1B, dotted arcs): The floor was rotated counterclockwise by 22.5° and the CS+ paired with sucrose and the CS-without sucrose were presented for 7 minutes.
3. Pause (Figure 1B, dashed arcs): The floor was rotated by 22.5° and replaced by a new floor without odorants or sucrose. The pause lasted for 5 seconds.
4. Memory test: The floor was rotated by 22.5° and replaced by a floor which had the CS+ and CS-but without sucrose. CS+ and CS-positions were switched from conditioning. The test phase lasted for 7 minutes.

Videos were recorded during the conditioning and test. After each experimental run, all flies were discarded and the Teflon fabric floor was rinsed with hot water and soap (Buzil G 530) using a sponge and dried over night to remove the odorants and the sucrose patch.

### Fly tracking

Video recordings were analyzed using the software Fiji (ImageJ 1.51s Wayne Rasband NIH, USA). We removed the first 30 frames due to compression artifacts and converted the video to grayscale. Then we did a Z projection to get the maximum intensity projection over the whole video, and calculated the difference per frame between the maximum intensity projection and the original video. This gave us a clear image of flies moving around the arena for tracking. We used this output to track flies using the plugin TrackMate (Version: 3.5.3, Tinevez et al., (2017)). We used a Downsample LoG detector to identify flies (blob diameter = 13 pixels, downsampling = 3, threshold = 6-8). To generate the tracks, we used the Simple LAP Tracker, with the following parameters: linking distance = 150 pixels, max gap closing = 150 pixels and maximal frame gap = 3 frames. For the conditioning data, we only extracted the x and y coordinates of each fly per frame. For the test data, we extracted the x and y coordinates per frame as well as the identity of the fly throughout the recording. We inspected all tracking results visually and corrected the tracks manually to connect the missing links and afterwards we extracted the x and y coordinates for the analysis.

### Data analysis

#### Normalizing arenas for comparison

For both the conditioning and the test datasets, we centralized each arena so that the center point of the circular arena was at (0, 0). The center point was determined by taking the midpoint between the CS+ and CS− locations; the x and y coordinates of the CS+ and CS− were recorded manually. We then converted each Cartesian coordinate to polar coordinates, in order to rotate each arena so that the CS+ location was at the top of the arena and the CS− was at the bottom. We took the distance of the CS+ to the center as a reference radius of 1, and normalized all coordinates to this radius. We then filtered out any points that had a radius equal to or greater than 1.3 to remove tracking errors. Note that for the conditioning dataset, only the x and y coordinates of a fly per frame were recorded. For the test dataset, the x and y coordinates per frame were recorded, but also the identity of the fly across frames.

#### Visit probability maps

Visit probability maps were generated only for the test dataset. For every individual fly, we divided the arena into 20 × 20 pixel bins. For each frame, we gave the pixel bin that contained the coordinate of the fly a score of 1, and gave all of the other bins a score of 0. We summed the scores of each pixel bin over all frames and then normalized by the number of frames that the individual was tracked for. For each group size, we then took the mean of each pixel bin over all individual fly tracks. We used the same analysis for the time-binned visit probability maps by looking only at the frames that occurred during the time bin.

#### Distance to the CS+ and CS−

We calculated the distance of each fly to the CS+ and to the CS-in every frame using:

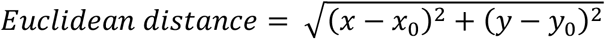

Where x and y are the Cartesian coordinates of the fly, and x_0_ and y_0_ are the Cartesian coordinates of either the CS+ or the CS−.

#### Relative distance to the CS+

For both the conditioning and test data, we calculated the relative distance to the odorant-sucrose patch (CS+ during test) of each fly per frame using:

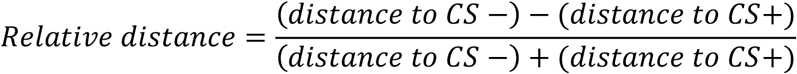

We then took the mean relative distance for the entire experimental run and for 1-minute time bins.

#### Preference index/Conditioned preference index

For both the conditioning and memory test data, the coordinates of the flies were tracked for each frame. For every frame of the experiment, we counted the number of flies in the arena half containing the odorant (CS+)-sucrose patch (during conditioning) or the CS+ (during memory test) and in the arena half containing the unrewarded odorant (CS-). We then calculated a preference index for each frame using:

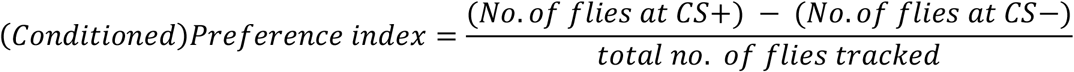

“Preference index” refers to the conditioning phase and “Conditioned preference index” refers to the memory test phase. We calculated the mean index for the entire experimental run and for 1-minute time bins. A preference index of 0 indicates that an equal number of flies were at the CS+ and the CS−, whereas a value of 1 indicates that all of the flies were in the half of the arena containing the CS+ and a value of 0 indicates that all of the flies were in the half of the arena containing the CS−.

#### Relative latency to the CS+

For the memory test dataset, we identified for each individual the time of the first frame that the individual was 0.5 cm or closer to the center of the CS+ and the center of the CS-. We then calculated the relative latency to reach the CS+ using:

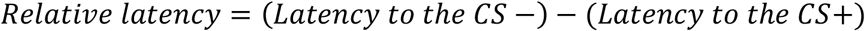

We calculated the mean relative latency per experimental run.

#### Distance between flies

For the memory test dataset, we used the rotated Cartesian coordinates to calculate the Euclidean distance between every fly in each frame of the experiment. We divided the distances into bins of 5 mm and counted the occurrences of each distance per experimental run.

#### Simulating distances between flies due to chance

For the test dataset, we selected flies according to their group size, and randomly sampled entire fly tracks from different experimental runs. We simulated as many experimental runs as there were real experimental runs, and we also simulated as many flies as were in each experimental run. We then overlaid these tracks and calculated the Euclidean distance between flies for the simulated experiments as we did for the real experiments.

#### Encounters between flies

We defined an encounter as the center of one fly being maximum two fly lengths (5 mm) away from the center of another fly. We calculated the Euclidean distance between all flies as before. We selected the distances that were less than or equal to 5 mm (the encounter distances). Since we had the identity of every fly per experimental run, we could calculate the number of encounters for every fly and the length of these encounters.

For the mean encounter number per fly per experimental run, we calculated the total number of encounters for one experimental run, multiplied it by two as there were two flies involved in each encounter, and then divided it by the number of flies in the arena. For the mean encounter length per experimental run, we summed the length of all encounters and divided by the total number of encounters per experimental run. We repeated this analysis for the simulated data.

### Statistical analysis

For all data analysis, R version 3.5.0 was used (R Core Team, 2018). All statistics were performed using Bayesian data analysis, based on Korner-Nievergelt et al., (2015). We chose Bayesian analysis over frequentist statistics as it allows us to a) estimate the probability with which the means of two groups differ, and b) determine the 95 % credible intervals within which the true mean of a group lies. Note that the frequentist confidence interval does not allow such a straightforward interpretation.

To investigate the effect of group size on behavioral performance (preference index (Fig. 1D, 2A, 2B), distance to the CS+ (Fig. S1A, S1B, S1C, S1D) and relative distance (Fig. S1E, S1F, S1G, S1H), we fitted a linear model (LM). The group size (“single”, “pair”, “small group” and “large group”) was used as the explanatory variable, with the large group as the reference level. The mean value per experimental run was used as the response variable. We used an improper prior distribution (flat prior) and simulated 100 000 values from the posterior distribution of the model parameters using the function “sim” from the package “arm”. The means of the simulated values from the posterior distributions of the model parameters were used as estimates, and the 2.5 % and 97.5 % quantiles as the lower and upper limits of the 95 % credible intervals. We used this linear model to compare the values of each group size against chance. For each group size, we calculated the proportion of simulated values from the posterior distribution that were larger than 0. If the proportion of simulated values was greater than 0, flies preferred the arena half containing the CS+ over the arena half containing the CS− (represented by filled circles in plots).

**Figure 2:**
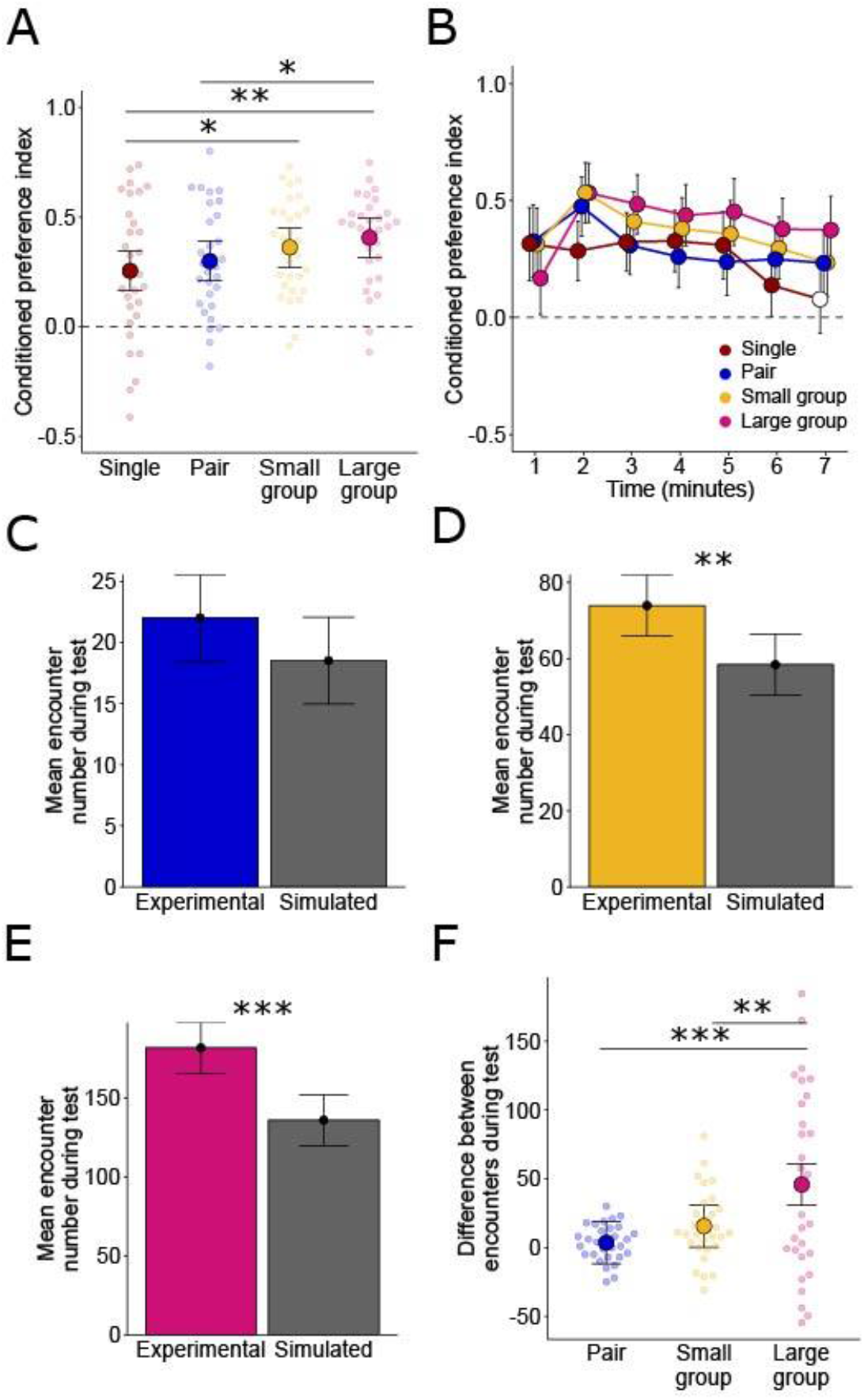
Group size affects associative odor-food memory expression and inter-fly encounters during the memory test. (**A**) Conditioned preference index for the different group sizes during the test. Large points represent the mean conditioned preference index per group size. Whiskers represent the 95 % credible intervals. Small points represent the mean conditioned preference index per experimental run. The dashed line indicates the conditioned preference index due to chance (0). Stars represent differences with Bayesian probabilities equal to or greater than 0.95 (*) or 0.99 (**) (N= 30 experimental runs). (**B**) Same data as in (A) but conditioned preference index over one-minute time bins during the test. Colors, dashed line and whiskers are the same as in (A). The points represent the mean conditioned preference index across all experimental runs within a time bin. Filled points are conditioned preference scores that are significantly different to chance. All groups preferred the arena half containing the CS+ patch than the arena half containing the CS− during all time bins (p(preference index > 0) ≥ 0.967), except for the single flies during the 7^th^ minute. For statistical comparisons between groups, see Table S1. (**C**) Mean inter-fly encounter number per fly per experimental run for the pair (blue) and the simulated pair (grey). Bars represent the mean across experimental runs (N= 30 experimental runs). Vertical lines represent the 95 % credible intervals. (**D**) Same as (C) for the small group (yellow) and the simulated small group (grey). (**E**) Same as (C) for the large group (pink) and the simulated large group (grey). Stars represent differences with Bayesian probabilities equal or greater 0.99 (**) or 0.999 (***) between the real and simulated group. (**F**) Paired differences between mean encounter numbers across experimental runs (N= 30 experimental runs). Large points represent the mean difference in encounter number between experimental runs. Small points represent the difference between the mean encounter numbers for each randomly assigned pair of real and simulated experimental runs. Colors and whiskers are the same as in A.

To test for differences between different group sizes, we calculated the proportion of simulated values from the posterior distribution that were larger for one group compared to another group. We declared an effect to be significant if the proportion was greater than or equal to 0.95 (*). Proportions greater than or equal to 0.99 are marked “**” and greater than or equal to 0.999 marked “***”. We performed this analysis for the whole recording: for the different time bins, and we compared the preference index and conditioned preference index of the different group sizes within a single time bin, not between them.

To test whether the relative latency (Fig. S1I) depends on group size, we used a LM as before. The relative latency was the response variable, and the group size was used as the explanatory variable. We used the same methodology as previously to simulate values from the posterior distribution and generate the means and the 95 % credible intervals. To test for differences, we calculated the proportion of draws from the posterior distribution for which the mean of each draw was smaller in the experimental dataset than the mean of each draw of the simulated dataset.

#### Inter-fly encounters

To investigate whether grouped flies differ in the number of their inter-fly encounters from random (simulated data), we used a LM for each distance bin. The number of occurrences of that distance was the response variable, and the type of data (experimental or simulated data) was used as the explanatory variable. We used the same method as specified above to test for differences.

To investigate whether the encounter number and lengths were different to random (simulated data), we used an LM with either encounter number or encounter length as the response variable, and the type of data (experimental or simulated data) as the explanatory variables. We used the same method as specified above to test for differences.

To determine whether the mean encounter number per fly differed between group sizes, we randomly assigned pairs of experimental and simulated encounter numbers from different experimental runs for each group size (Figure 2F). For each pair, we then subtracted the simulated encounter number value from the real encounter number value (difference between encounters). This allowed us to compare between group sizes as by removing the simulated value, we remove the number of encounters that could be due to chance, which is positively correlated with group size. To test for differences between group sizes, we used a LM. The response variable was the “difference between encounters”. The explanatory variable was the different group size (“pair”, “small group” and “large group”). The large group was used as the reference level. We used the same method as specified above to draw inferences about the differences between the large group and the other two group sizes.

## RESULTS

To investigate whether group size affects associative odor-food memory in flies, we developed an automated assay to condition four groups of flies simultaneously (Figure 1A, B). The learning assay consisted of a computer-controlled rotating platform with four circular arenas. This design allowed us to transfer flies from one experimental phase to another without anesthesia and with minimal mechanical disturbance, which can alter fruit flies’ behavior (Barron, 2000; Bartholomew et al., 2015; Trannoy et al., 2015). We compared four different sized groups of flies: one fly (single), two flies (pair), three or four flies (small group) and seven or eight flies (large group) (Figure 1C). Both the training phase and test phase lasted for 7 minutes. During the 7-minutes long conditioning, two odorants (2,3-butanedione and ethyl acetate) were presented at opposite sides of the arena; one odorant (conditioned stimulus, CS+) was paired with dried sucrose and the other odorant was not (CS−). Each odorant was used equally often as CS+ and CS− and the data were pooled. This procedure minimizes non-associative effects of the conditioning, such as odorant-specific changes in hedonic value, generalization or sensitization (Quinn et al., 1974). During conditioning, flies were allowed to forage around the arena and find the food and odorant source.

### Flies aggregate on the odorant-sucrose patch during conditioning and learn to associate the odorant with sucrose

To determine whether flies approached the odorant-sucrose patch, we counted the number of flies in the half of the arena containing the odorant-sucrose patch, in order to calculate a mean preference index for each experimental run (see Materials and Methods). Flies of all group sizes showed a higher preference for the arena half containing the odorant (CS+)−sucrose patch than for the arena half containing the CS− (Figure 1D). From the second minute to the end of the conditioning, the probability that flies preferred the arena half containing the odorant (CS+)−sucrose patch over the arena half containing the CS− was above 0.999 across all groups, implying that they were feeding on the sucrose (see Table S1 for the Bayesian probabilities comparing between groups). To confirm that the preference index reliably measures flies’ preference, we additionally calculated the mean distance to the CS+ per experimental run (Figure S1A, S1B and Table S1) and the distance to the CS+ relative to the distance to the CS− (relative distance, Figure S1E, S1F and Table S1). Both measures revealed similar behaviors as the preference index, showing that flies of all group sizes approached the CS+ and remained within 10 mm of its center from the 4^th^ minute onwards (Figure S1B). Flies in pairs remained at larger distance from the CS+ than single flies or flies in the small group (Figure S1A, S1B, S1E, S1F). A possible explanation for the larger distance from the CS+ in pairs could be a higher aggression rate, as aggression between female flies depends on group size and flies exhibit less aggression when kept in groups as compared to kept in isolation (Ueda and Kidokoro, 2002).

Flies were transferred from the conditioning to the test by rotating the platform (Figure 1B). In between conditioning and test there was a pause, where the platform was rotated to a neutral segment where there were no odorants or sucrose present. The pause lasted 5 s, and then platform was rotated to the test segment. During the test, the positions of the CS+ and the CS− were switched and there was no sucrose, thus flies could not rely on remembering the location of the sucrose patch (Kim and Dickinson, 2017) and had to follow the olfactory CS+ to search for the expected food. To visualize the conditioned preference index during the test, we projected the flies’ trajectories on a plane and calculated the probability across flies to visit a particular pixel bin (Figure 1E). The increased visit probabilities around the CS+ persisted over the entire 7 minutes of the test, confirming that flies learned to associate the CS+ with food (Figure 1E).

### Associative odor-food memory expression increases with group size

We next asked whether group size affects the expression of associative odor-food memory during the test and determined the conditioned preference index for the CS+ (Figure 2A). During the test, flies of all group sizes preferred the arena half containing the CS+ over the arena half containing the CS− (*p(all group sizes > 0)* > 0.999), showing that flies had formed an associative odor-food memory (Figure 2A). Flies of the large group showed a higher conditioned preference for the CS+ than the pair and the single fly (*p(large group > pair)* = 0.95, *p(large group > single)* = 0.99), and the small group showed a higher conditioned preference than the single fly *(p(small group > single)* = 0.95) (see Table S1). Flies showed a similar group size-dependence of their conditioned responses when we measured the distance to the CS+ (Figure S1C, S1D and Table S1) or the relative distance to the CS+ (Figure S1G, S1H, Table S1). There were no differences between groups in the latency to arrive at the CS+ relative to the latency to arrive at the CS− (Figure S1I).

To investigate the time course of memory expression, we calculated the conditioned preference index over one-minute time bins (Figure 2B). During the first minute, there were no differences of the conditioned preference between any of the group sizes (Table S1). Between the 2^nd^ and 7^th^ minutes, flies from the large group showed higher conditioned preference than single or paired flies throughout most bins tested (Table S1). In all minute bins, the conditioned preference was higher than chance for all group sizes, except for the single flies in the 7^th^ minute (p(preference index > 0) = 0.861). The distance to the CS+ (Figure S1D, Table S1) and the relative distance (Figure S1H, Table S1) revealed similar group size-dependent differences in the conditioned approach behavior. These results suggest that the expression of an associative odor-food memory increases in strength and duration with increasing group size.

### Flies tested in groups – but not in pairs – exhibit more inter-fly encounters than random

The extended associative memory expression in flies of the small and large group indicates that flies socially interact to share information about the location of the predicted food source, as has previously been found during foraging (Abu et al., 2018; Lihoreau et al., 2016; Tinette et al., 2004). If the extended memory expression in the small and large group depends on social interactions, then the group size should affect the frequency of social interactions. To assess whether group size affects the frequency of social interactions we measured the number of inter-fly distances and compared it with the number of expected random distances. We calculated the distances between all flies conditioned in groups in each video frame during conditioning to see if they approached each other. To determine whether these distance distributions could be explained due to flies randomly encountering each other in the arena, we simulated 30 new experimental runs by randomly sampling fly locations from all experimental runs for each video frame. We then calculated the distances between these simulated groups of flies in each video frame (Figure S2A). For the large group there were more short inter-fly distances (0-5 and 5-10 mm) than for the simulated group.

We chose a distance of 5 mm between fly centers as a threshold for inter-fly encounters where flies could potentially socially interact. Flies in the small and large group made more encounters (approached each other by 5 mm or less) than the simulated groups of flies, but not the pair (*p(large group > simulated large group)* > 0.999, *p(small group > simulated small group)* = 0.996, *p(pair > simulated pair)* = 0.917) (Figure 2C-E).

To compare encounter number across group sizes, we needed to correct for trivial differences in encounters that are just due to differences in the group sizes (in larger groups there is a higher chance for random inter-fly encounters). We corrected for these differences in encounter number by the following procedure: We randomly took an experimental run from the experimental and simulated datasets and subtracted the number of encounters between the two experimental runs (Figure 2F). By subtracting the number of encounters in the simulated runs, we removed the number of encounters per experimental run that could be due to random encounters. The encounter number was higher for the large group compared to the small group and the pair *(p(large group > pair)* > 0.999, *p(large group > small group)* = 0.996). There were no differences in encounter length between any group size and their simulated groups (Figure S2B).

The increased number of encounters in the larger group indicates that flies are more attracted to each other when they are in large groups than when they are in small groups or pairs. More encounters allow more opportunities for social interactions between flies, which in turn could underlie the longer associative memory expression of the large group as compared to smaller groups or single flies.

## DISCUSSION

We developed an automated learning and memory assay for walking fruit flies that allows analyzing the behavior of individual flies while they forage, learn and memorize odor-food associations alone or in groups. The strength and duration of odor-food memory expression increased with group size, and flies in larger groups were more attracted to each other than flies in smaller groups. These data suggest that foraging in groups facilitates social information transfer about the quality of a food source (during odor-food learning) or about the location of the predicted food source (during olfactory memory-guided food search).

### Social information transfer during foraging

Fruit flies accumulate on fermenting fruit which they find by following both odorants released by fermenting fruit (Becher et al., 2012; Kellogg et al., 1962; Semmelhack and Wang, 2009) and aggregation pheromones released by male (Bartelt et al., 1985; Lin et al., 2015; Mercier et al., 2018) and female conspecifics (Lebreton et al., 2017). During foraging, primer flies explore the environment and appear to signal the location of favorable food patches to other flies (Tinette et al., 2004). Besides sharing information about food sources, fruit flies also share information about their internal state, such as stress (Suh et al., 2004), and about the location and quality of resources during mate choice (Danchin et al., 2018; Mery et al., 2009) and egg-laying sites (Battesti et al., 2012; Durisko and Dukas, 2013; Lin et al., 2015; Sarin and Dukas, 2009).

Our finding of extended expression of associative odor-food memories in groups, together with the positive correlation between group size and inter-fly attraction, suggests that associative odor-food learning or memory expression also benefits from social information transfer during aggregation. The positive correlation between group size and inter-fly attraction that we found is in line with a previous study where inter-fly attraction was higher in larger than in smaller groups (20-40 versus 10 flies) (Simon et al., 2012). To our knowledge, such an increase in inter-animal attraction with increasing group size has not yet been reported in vertebrates (Miller and Stephen, 1966).

### Social effects on odor-food learning

The increased associative odor-food memory with increasing group size could be a result of social information transfer during the learning of the odor-food association (during conditioning) or during the retrieval of the odor-food memory (during the memory test). Since our experiments were performed in the dark, flies could have transferred information socially via olfactory stimuli (Jallon, 1984; Keesey et al., 2016; Lebreton et al., 2017; Lin et al., 2015), gustatory stimuli (Schneider et al., 2012) sound (Tauber and Eberl, 2003), substrate-borne vibration (Fabre et al., 2012) and touch (Ramdya et al., 2014), but not via visual cues (Danchin et al., 2018; Ferreira and Moita, 2019; Golden and Dukas, 2014; Kim et al., 2012; Mery et al., 2009; Sarin and Dukas, 2009).

During conditioning, the presence of other flies at the sucrose patch could increase the reinforcing strength of the sucrose since the presence of other flies indicates that the food patch is good. Indeed, flies prefer food sources with other flies present over food sources without any flies (Lihoreau et al., 2016; Tinette et al., 2004). Alternatively, the presence of other flies at the sucrose patch may be an additional appetitive reinforcing stimulus. The role of conspecifics as a positive reinforcer has been previously demonstrated in honey bees, where antennal touching of a nestmate acts as positive reinforcer during odor conditioning (Cholé et al., 2019). The fact that fruit flies are attracted to each other (Lefranc et al., 2001; Simon et al., 2012; Tinette et al., 2004) makes it plausible to assume that flies could also act as an additional positive reinforcer at the sucrose patch.

The reinforcing function of other flies could be mediated by dopaminergic neurons, because dopaminergic neurons mediate the reinforcing function of sucrose (Liu et al., 2012) and because dopamine itself has an effect on the sociality of flies: inter-fly attraction decreases in flies that have a deficiency in dopamine released from neurons and hypodermal cells (Fernandez et al., 2017).

### Social effects on odor-food memory expression

Besides a learning effect, the extended odor-food memory expression could be a memory retrieval effect. Memory retrieval could be affected by the social interactions during the memory test, as flies that had learned the association between CS+ and sucrose could transfer information about the location of the predicted sucrose patch to flies that have failed to learn. Information transfer from experienced to naïve flies can affect group level behavior during odor avoidance (Ramdya et al., 2014), aversive memory retrieval (Chabaud et al., 2009), mate choice (Danchin et al., 2018; Mery et al., 2009), oviposition site choice (Battesti et al., 2012; Sarin and Dukas, 2009) and predator-induced egg-retention (Kacsoh et al., 2015). Moreover, a theoretical study predicted that social interactions can increase performance during odor-guided foraging (Torney et al., 2009).

Alternatively, flies that located the CS+ first during the test could serve as an attractive reinforcing stimulus (see discussion above and (Lihoreau et al., 2016; Tinette et al., 2004)), thus appetitive learning of the CS+ could continue throughout the test. This ongoing appetitive learning of the CS+ during the test would appear as extended associative memory expression in our study.

Another possible explanation for the extended memory expression could be reduced memory extinction due to social interactions at the location of the CS+. Memories can be extinguished when the CS+ is presented without reinforcement (Lagasse et al., 2009; Schwaerzel et al., 2002), which is effectively what occurs throughout the test in our study. In flies, extinction of odor-sucrose memories is mediated by dopaminergic neurons that encode punishment (Felsenberg et al., 2017): lack of reward during CS+ induced memory retrieval activates punishment-encoding dopaminergic neurons, and this activation counteracts the associative odor-food memory. The presence of other flies at the CS+ could provide an appetitive stimulus and thereby prevent the activation of these extinction mediating dopaminergic neurons.

### Limitations of the study and outlook

We found a positive relationship between group size and the strength and duration of odor-food memory expression. However, our experimental design does not allow conclusions on whether this extended memory expression results from being in the group during odor-food learning (conditioning) or during olfactory memory-guided search (memory test). To discriminate between these two possibilities one could test whether flies conditioned in a group and tested alone (or conditioned alone and tested in a group) still show extended memory expression as compared to control flies that were conditioned and tested alone.

We analyzed the walking behavior of individual flies during the memory test, but not during the conditioning because we could not separate flies from each other when they clustered at the sucrose patch due to a lack of spatial resolution. By using cameras with higher spatial resolution, this assay can be extended to a high-throughput assay for tracking individuals in multiple parallel fly groups, allowing classification of pairwise and higher-order interactions between individuals, as well as stereotyped behaviors in individuals (Berman et al., 2016; Branson et al., 2009; Onodera et al., 2019).

This assay could help reveal external factors (e.g., fly density, the ratio of informed to uninformed flies) and internal factors (e.g., sex, metabolic, genetic, or circadian states) that influence learning and memory expression in social contexts. Importantly, this assay would allow studying the neural basis of social effects on foraging, by disentangling sensory processing and memory formation. To identify the sensory bases of information transmission between flies, one could test the effect of temporarily perturbing their ability to smell, see and mechanosense by expressing a temperature-sensitive switch for synaptic transmission in defined neuron populations (Kim et al., 2012; Kitamoto, 2001; Ramdya et al., 2014). Likewise, neuronal perturbation experiments would help identifying the neurons that encode the valence of social information and reveal how these neurons integrate with the neurons known to encode the hedonic and caloric value of food (Huetteroth et al., 2015). Moreover, to investigate whether information transmission during foraging is affected by the fly’s predisposition to forage, one could use the two naturally occurring *foraging* gene *Drosophila* mutants. “Rovers” move more during foraging and demonstrate improved short term memory, whereas “sitters” move less and show an improved long term memory (Mery et al., 2007; Osborne et al., 1997). Since both foraging and aversive memory expression are affected by social context (Kohn et al., 2013), experiments using these morphs would help to assess the genetic bases of social effects on odor-food learning and memory expression.

## ACKNOWLEDGEMENTS

We thank C. Giovanni Galizia for discussions initiating this research, Stefanie Neupert for programming the acquisition software and comments on the manuscript, Jana Hörsch for contributing to pilot experiments and comments on the manuscript, Charles Ellen for comments on the manuscript, FIBERFLON (Konstanz) for donating Teflon-coated fiberglass fabric.

## AUTHOR CONTRIBUTIONS

PS conceptualized and designed the study. CT performed the data collection. CT and YM performed the pilot experiments. AS, CT and YM prepared the video data for analysis. AS performed the statistical analysis. AS, PS and YM wrote the manuscript. PS supervised the study.

## SUPPLEMENTARY MATERIALS

**Table S1:**
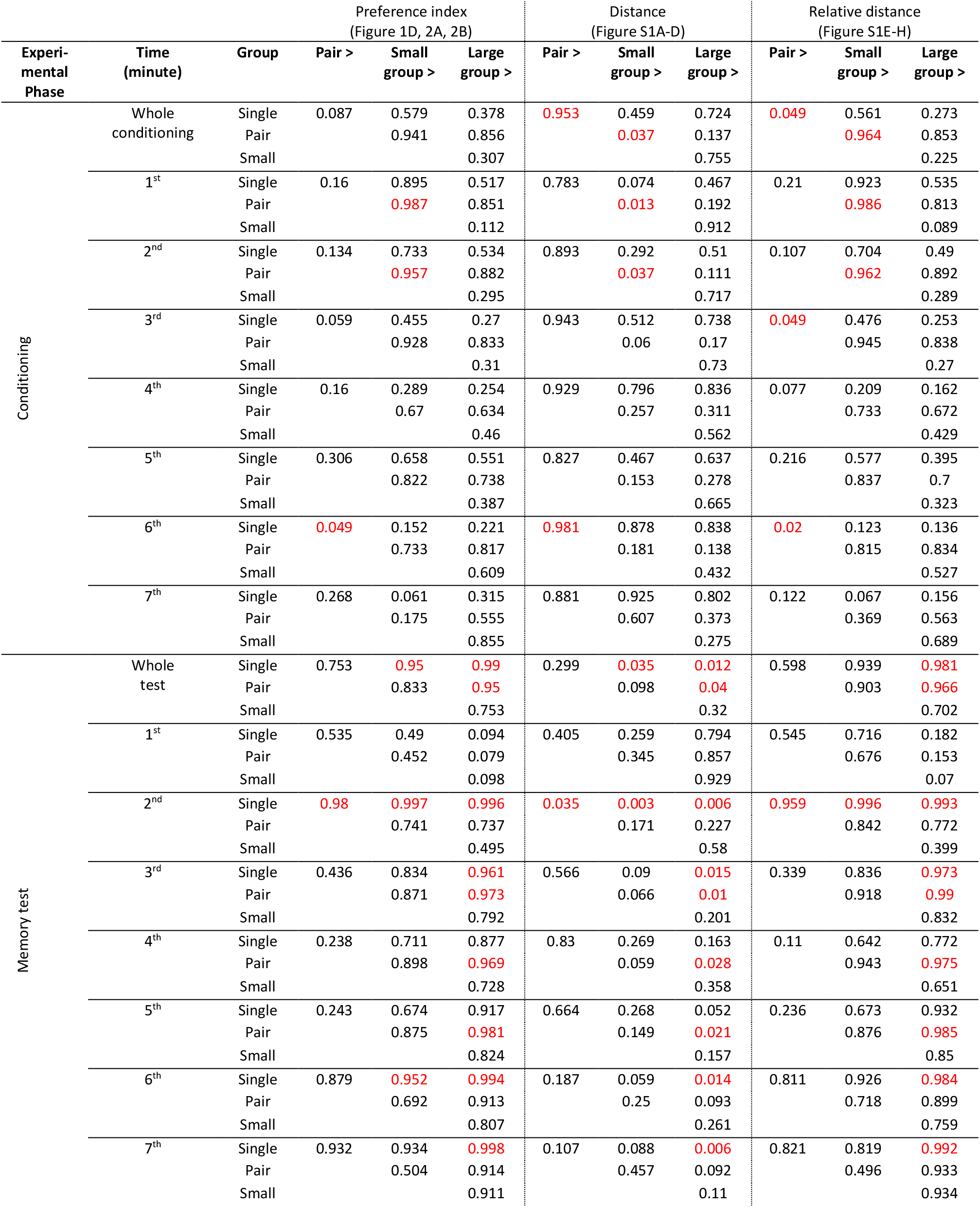
Bayesian probabilities comparing the preference index, distance and relative distance between groups. Related to Figure 1D, 2A, 2B and S1. Bayesian probability with which the mean of a group in the column “Pair >”, “Small group >” or “Large group >” is larger than the mean of a group in the column “Group”. Red values indicate probabilities equal to or greater than 0.95 or equal to or smaller than 0.05.

| Experi-<br>mental<br>Phase | Time<br>(minute) | Group | Preference index<br>(Figure 1D, 2A, 2B) |  |  | Distance<br>(Figure S1A-D) |  |  | Relative distance<br>(Figure S1E-H) |  |  |
| --- | --- | --- | --- | --- | --- | --- | --- | --- | --- | --- | --- |
|  |  |  | Pair > | Small<br>group > | Large<br>group > | Pair > | Small<br>group > | Large<br>group > | Pair > | Small<br>group > | Large<br>group > |
| Conditioning | Whole<br>conditioning | Single | 0.087 | 0.579 | 0.378 | 0.953 | 0.459 | 0.724 | 0.049 | 0.561 | 0.273 |
|  |  | Pair |  | 0.941 | 0.856 |  | 0.037 | 0.137 |  | 0.964 | 0.853 |
|  |  | Small |  |  | 0.307 |  |  | 0.755 |  |  | 0.225 |
|  | 1<sup>st</sup> | Single | 0.16 | 0.895 | 0.517 | 0.783 | 0.074 | 0.467 | 0.21 | 0.923 | 0.535 |
|  |  | Pair |  | 0.987 | 0.851 |  | 0.013 | 0.192 |  | 0.986 | 0.813 |
|  |  | Small |  |  | 0.112 |  |  | 0.912 |  |  | 0.089 |
|  | 2<sup>nd</sup> | Single | 0.134 | 0.733 | 0.534 | 0.893 | 0.292 | 0.51 | 0.107 | 0.704 | 0.49 |
|  |  | Pair |  | 0.957 | 0.882 |  | 0.037 | 0.111 |  | 0.962 | 0.892 |
|  |  | Small |  |  | 0.295 |  |  | 0.717 |  |  | 0.289 |
|  | 3<sup>rd</sup> | Single | 0.059 | 0.455 | 0.27 | 0.943 | 0.512 | 0.738 | 0.049 | 0.476 | 0.253 |
|  |  | Pair |  | 0.928 | 0.833 |  | 0.06 | 0.17 |  | 0.945 | 0.838 |
|  |  | Small |  |  | 0.31 |  |  | 0.73 |  |  | 0.27 |
|  | 4<sup>th</sup> | Single | 0.16 | 0.289 | 0.254 | 0.929 | 0.796 | 0.836 | 0.077 | 0.209 | 0.162 |
|  |  | Pair |  | 0.67 | 0.634 |  | 0.257 | 0.311 |  | 0.733 | 0.672 |
|  |  | Small |  |  | 0.46 |  |  | 0.562 |  |  | 0.429 |
|  | 5<sup>th</sup> | Single | 0.306 | 0.658 | 0.551 | 0.827 | 0.467 | 0.637 | 0.216 | 0.577 | 0.395 |
|  |  | Pair |  | 0.822 | 0.738 |  | 0.153 | 0.278 |  | 0.837 | 0.7 |
|  |  | Small |  |  | 0.387 |  |  | 0.665 |  |  | 0.323 |
|  | 6<sup>th</sup> | Single | 0.049 | 0.152 | 0.221 | 0.981 | 0.878 | 0.838 | 0.02 | 0.123 | 0.136 |
|  |  | Pair |  | 0.733 | 0.817 |  | 0.181 | 0.138 |  | 0.815 | 0.834 |
|  |  | Small |  |  | 0.609 |  |  | 0.432 |  |  | 0.527 |
|  | 7<sup>th</sup> | Single | 0.268 | 0.061 | 0.315 | 0.881 | 0.925 | 0.802 | 0.122 | 0.067 | 0.156 |
|  |  | Pair |  | 0.175 | 0.555 |  | 0.607 | 0.373 |  | 0.369 | 0.563 |
|  |  | Small |  |  | 0.855 |  |  | 0.275 |  |  | 0.689 |
| Memory test | Whole<br>test | Single | 0.753 | 0.95 | 0.99 | 0.299 | 0.035 | 0.012 | 0.598 | 0.939 | 0.981 |
|  |  | Pair |  | 0.833 | 0.95 |  | 0.098 | 0.04 |  | 0.903 | 0.966 |
|  |  | Small |  |  | 0.753 |  |  | 0.32 |  |  | 0.702 |
|  | 1<sup>st</sup> | Single | 0.535 | 0.49 | 0.094 | 0.405 | 0.259 | 0.794 | 0.545 | 0.716 | 0.182 |
|  |  | Pair |  | 0.452 | 0.079 |  | 0.345 | 0.857 |  | 0.676 | 0.153 |
|  |  | Small |  |  | 0.098 |  |  | 0.929 |  |  | 0.07 |
|  | 2<sup>nd</sup> | Single | 0.98 | 0.997 | 0.996 | 0.035 | 0.003 | 0.006 | 0.959 | 0.996 | 0.993 |
|  |  | Pair |  | 0.741 | 0.737 |  | 0.171 | 0.227 |  | 0.842 | 0.772 |
|  |  | Small |  |  | 0.495 |  |  | 0.58 |  |  | 0.399 |
|  | 3<sup>rd</sup> | Single | 0.436 | 0.834 | 0.961 | 0.566 | 0.09 | 0.015 | 0.339 | 0.836 | 0.973 |
|  |  | Pair |  | 0.871 | 0.973 |  | 0.066 | 0.01 |  | 0.918 | 0.99 |
|  |  | Small |  |  | 0.792 |  |  | 0.201 |  |  | 0.832 |
|  | 4<sup>th</sup> | Single | 0.238 | 0.711 | 0.877 | 0.83 | 0.269 | 0.163 | 0.11 | 0.642 | 0.772 |
|  |  | Pair |  | 0.898 | 0.969 |  | 0.059 | 0.028 |  | 0.943 | 0.975 |
|  |  | Small |  |  | 0.728 |  |  | 0.358 |  |  | 0.651 |
|  | 5<sup>th</sup> | Single | 0.243 | 0.674 | 0.917 | 0.664 | 0.268 | 0.052 | 0.236 | 0.673 | 0.932 |
|  |  | Pair |  | 0.875 | 0.981 |  | 0.149 | 0.021 |  | 0.876 | 0.985 |
|  |  | Small |  |  | 0.824 |  |  | 0.157 |  |  | 0.85 |
|  | 6<sup>th</sup> | Single | 0.879 | 0.952 | 0.994 | 0.187 | 0.059 | 0.014 | 0.811 | 0.926 | 0.984 |
|  |  | Pair |  | 0.692 | 0.913 |  | 0.25 | 0.093 |  | 0.718 | 0.899 |
|  |  | Small |  |  | 0.807 |  |  | 0.261 |  |  | 0.759 |
|  | 7<sup>th</sup> | Single | 0.932 | 0.934 | 0.998 | 0.107 | 0.088 | 0.006 | 0.821 | 0.819 | 0.992 |
|  |  | Pair |  | 0.504 | 0.914 |  | 0.457 | 0.092 |  | 0.496 | 0.933 |
|  |  | Small |  |  | 0.911 |  |  | 0.11 |  |  | 0.934 |

**Figure S1:**
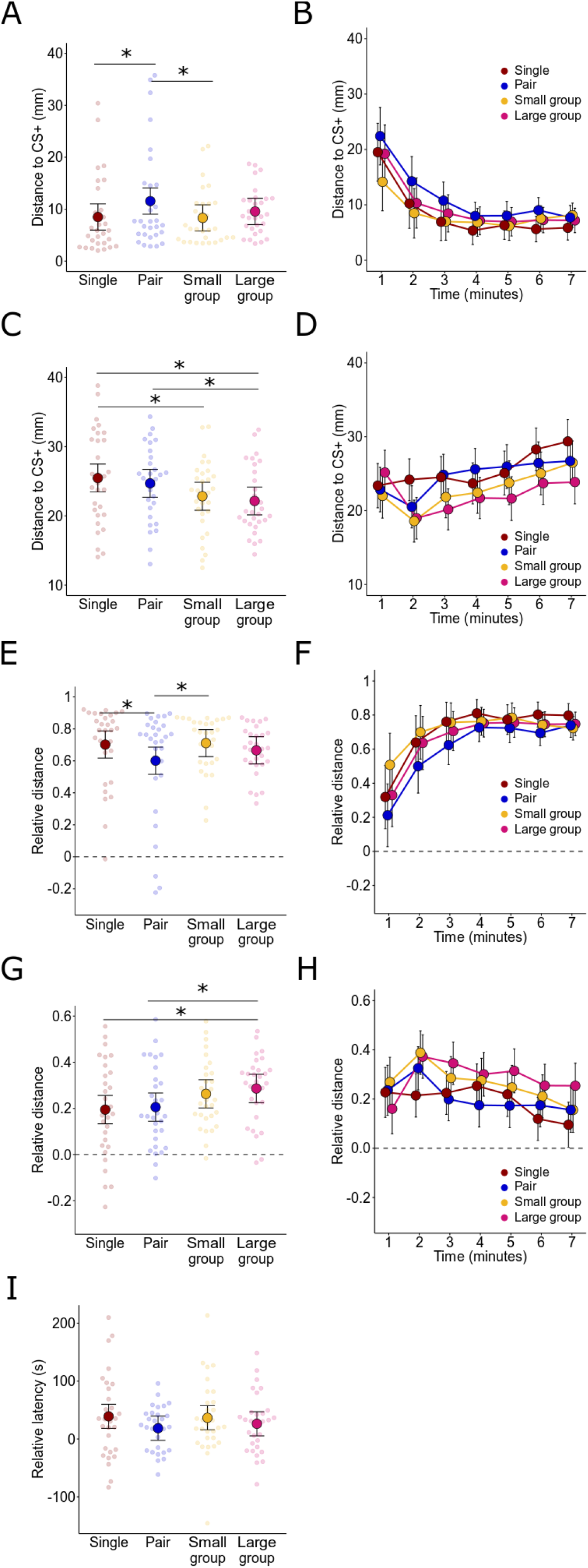
Flies’ behavioral performance during conditioning and memory test. Related to Figure 1D, 2A and 2B. (**A**) Distance to the CS+ of flies in the different groups during conditioning. Large points represent the mean distance to the CS+ per group size. Whiskers represent the 95 % credible intervals. Small points represent the mean distance to the CS+ per experimental run (N= 30 experimental runs). Stars represent differences with probabilities equal to or greater than 0.95. (**B**) Same data as in (A) but for one-minute time bins. Points represent the mean distance to the CS+ per group size. Colors and whiskers are the same as in (A). For statistical comparisons between groups, see Table S1 (applies to all figure panels). (**C**) Distance to the CS+ during the memory test. (**D**) Same data as in (C) but for one-minute time bins. (**E**) Relative distance of flies in the different groups during conditioning. The dashed line indicates the relative distance due to chance (0). (**F**) Same data as in (E) but for one-minute time bins. In all time bins, all groups were closer to the CS+-sucrose patch than to the CS− *(p(relative distance > 0)* ≥ 0.988). (**G**) Relative distance during the memory test. (**H**) Same data as in (G) but for one-minute time bins. In all time bins, all groups were closer to the CS+ than to the CS− *(p(relative distance > 0)* ≥ 0.98). (**I**) Relative latencies of flies to reach CS+ ([latency to the CS−] – [latency to the CS+]) for the different groups during the memory test. The Bayesian probabilities for intergroup-differences were below 0.92.

**Figure S2:**
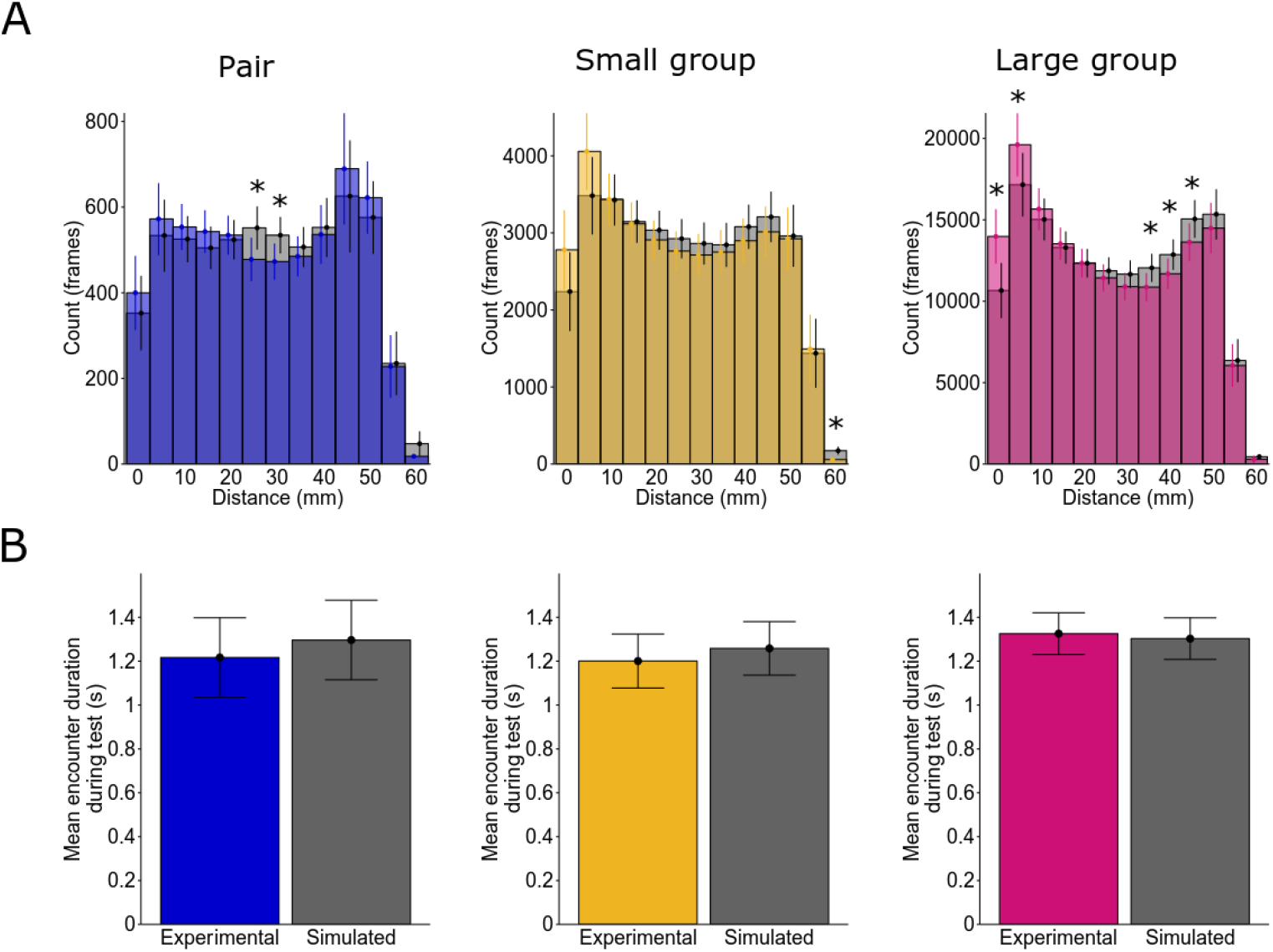
Inter-fly encounters during the test. Related to Figure 2C – E. (**A**) The number of inter-fly distances during the test for flies conditioned in pairs (left), small groups (middle) and large groups (right. The first bin ranges from 0 mm to 2.5 mm, and the following bins represent a range of 5 mm i.e. the 10 mm bin ranges from 7.5 mm to 12.5 mm. Bars represent the mean number of distances per bin between real groups of flies (pair: blue, small group: yellow, large group: pink) and between simulated groups of flies (grey). Vertical lines represent the 95 % credible intervals. Videos were recorded at 15 frames/s. Bars of the experimental and simulated number of distances within the same bin were compared to each other. Stars indicate differences with probabilities equal to or greater than 0.95 between the experimental and simulated data within a bin (N = 30 real or simulated experimental runs). (**B**) Mean encounter length per experimental run for the pair (blue, left), small group (yellow, middle) and large group (pink, right). The corresponding simulated groups of flies are shown in grey. Bars represent the mean across experimental runs. Whiskers represent the 95 % credible intervals. Stars represent differences with probabilities equal to or greater than 0.95 between the real and simulated groups.

